# Nanoparticle-delivered TLR4 and RIG-I agonists enhance immune response to SARS-CoV-2 subunit vaccine

**DOI:** 10.1101/2022.01.31.478507

**Authors:** Alexandra Atalis, Mark C Keenum, Bhawana Pandey, Alexander Beach, Pallab Pradhan, Casey Vantucci, Ritika Jain, Justin Hosten, Clinton Smith, Liana Kramer, Angela Jimenez, Miguel Armenta Ochoa, David Frey, Krishnendu Roy

## Abstract

Despite recent success in vaccinating populations against SARS-CoV-2, concerns about immunity duration, continued efficacy against emerging variants, protection from infection and transmission, and worldwide vaccine availability, remain. Although mRNA, pDNA, and viral-vector based vaccines are being administered, no protein subunit-based SARS-CoV-2 vaccine is approved. Molecular adjuvants targeting pathogen-recognition receptors (PRRs) on antigen-presenting cells (APCs) could improve and broaden the efficacy and durability of vaccine responses. Native SARS-CoV-2 infection stimulate various PRRs, including toll-like receptors (TLRs) and retinoic-acid-inducible gene I-like receptors (RIG-I). We hypothesized that targeting the same PRRs using adjuvants on nanoparticles along with a stabilized spike (S) protein antigen could provide broad and efficient immune responses. Formulations targeting TLR4 (MPLA), TLR7/8 (R848), TLR9 (CpG), and RIG-I (PUUC) delivered on degradable polymer-nanoparticles (NPs) were combined with the S1 subunit of S protein and assessed in vitro with isogeneic mixed lymphocyte reactions (iso-MLRs). For in vivo studies, the adjuvanted nanoparticles were combined with stabilized S protein and assessed using intranasal and intramuscular prime-boost vaccination models in mice. Combination NP-adjuvants targeting both TLR and RIG-I (MPLA+PUUC, CpG+PUUC, or R848+PUUC) differentially increased proinflammatory cytokine secretion (IL-1β, IL-12p70, IL-27, IFN-β) by APCs cultured in vitro, and induced differential T cell proliferation. When delivered intranasally, MPLA+PUUC NPs enhanced local CD4+CD44+ activated memory T cell responses while MPLA NPs increased anti-S-protein-specific IgG and IgA in the lung. Following intramuscular delivery, PUUC-carrying NPs induced strong humoral immune responses, characterized by increases in anti-S-protein IgG and neutralizing antibody titers and germinal center B cell populations (GL7+ and BCL6+ B cells). MPLA+PUUC NPs further boosted S-protein-neutralizing antibody titers and T follicular helper cell populations in draining lymph nodes. These results suggest that SARS-CoV-2-mimicking adjuvants and subunit vaccines could lead to robust and unique route-specific adaptive immune responses and may provide additional tools against the pandemic.

**GRAPHICAL ABSTRACT:** 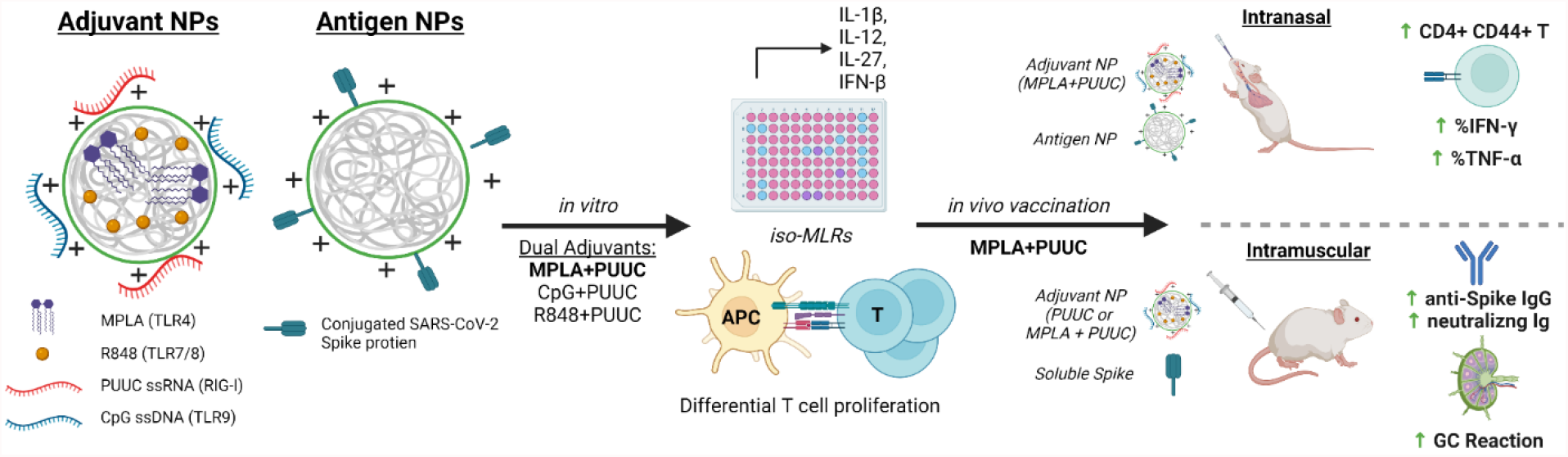

## INTRODUCTION

The coronavirus disease 2019 (COVID-19) pandemic caused by severe acute respiratory syndrome coronavirus 2 (SARS-CoV-2) has killed more than 5 million people worldwide.^1^ The crisis has elicited a global scientific effort to develop vaccines and therapies for the disease at an unprecedented speed. Phase I trials started as early as March 2020 and as of November 2021, there at least 322 vaccines in various stages of development.^2^ A non-replicating viral vector vaccine from Janssen and two mRNA-based vaccines from Moderna and Pfizer-BioNTech – all against SARS-CoV-2 spike (S) protein antigen – have been authorized for emergency use in the United States. The Pfizer-BioNTech vaccine received full FDA approval for individuals 16 and older in August 2021.^3^ However, much remains to be determined regarding the longevity of immune responses and correlates of protection against emerging SARS-CoV-2 variant strains in various patient populations. Recently, booster shots, 2 months after initial dose of the Janssen vaccine and 6 months after initial dose of the mRNA vaccines, were approved due to waning immunity amongst vaccinated individuals. Much work remains to develop long term immunity and to understand how both local immunity in the lung and the associated systemic immunity can protect against emerging variants in a durable manner.

The incorporation of molecular adjuvants in vaccines, especially protein subunit vaccines (which make up >35% of all vaccines in development), could be a potential strategy to induce more robust immune responses against SARS-CoV-2 through the targeting of receptors on antigen-presenting cells (APCs).^2,4^ Aluminum-containing adjuvants (e.g., alum) have been included in vaccines to enhance immunogenicity since the 1930s.^5^ Traditional adjuvants like alum have been developed and tested empirically, but in recent years vaccine design strategy has shifted to a more rational approach where each component elicits a defined immunological pathway to shift the immune response. Pathogen-associated molecular patterns (PAMPs), including lipids, carbohydrates, peptides, and nucleic acids commonly expressed by pathogens, are currently being investigated as adjuvants because they specifically bind to pattern-recognition receptors (PRRs) on APCs and induce maturation.^4^ Mature APCs, namely dendritic cells (DCs), initiate antigen-specific adaptive immune responses by activating naïve T cells, which differentiate into effectors, such as cytotoxic T cells (CD8+) and T helper cells (CD4+).^6^ T helper cells (type 2, Th2) mediate the differentiation of B cells into antibody-producing plasma cells.^7,8^

Two classes of PRRs are toll-like receptors (TLRs) and retinoic-acid-inducible gene I (RIG-I)-like receptors. RIG-I receptors are in the cytosol and recognize short double stranded RNA, a replication intermediate for RNA viruses, that exhibit viral motifs such as an uncapped 5’ diphosphate or triphosphate end.^9^ The poly-uridine core of Hepatitis C virus (HCV), poly-U/UC (PUUC), activates RIG-I and triggers potent anti-HCV responses.^10^ The most studied TLR-ligand combination is TLR4 and the Gram-negative bacterial cell wall component lipopolysaccharide (LPS), which includes monophosphoryl lipid A (MPLA), a molecule used in FDA-approved adjuvant systems (AS01 and AS04, GlaxoSmithKline).^11–13^ Endosomal TLRs 7 and 8 recognize single-stranded RNA and can be activated by synthetic imidazoquinolines including resiquimod (R848).^14^ Endosomal TLR9 recognizes single-stranded DNA, and current understanding suggests that TLR9 recognizes unmethylated CpG motifs common to bacteria and viruses to discriminate from self-DNA.^15^ Most TLRs (TLR4, TLR7/8, TLR9) signal through the MyD88 pathway, which activates NFkB to induce production of proinflammatory cytokines (i.e., IL-1β) and IRF7 to induce type I interferon secretion.^16–18^ When trafficked from the plasma membrane to endosomes, TLR4 can signal through the TRIF pathway and induce expression of IFN-β (**Figure 1A**).^18,19^

**Figure 1:**
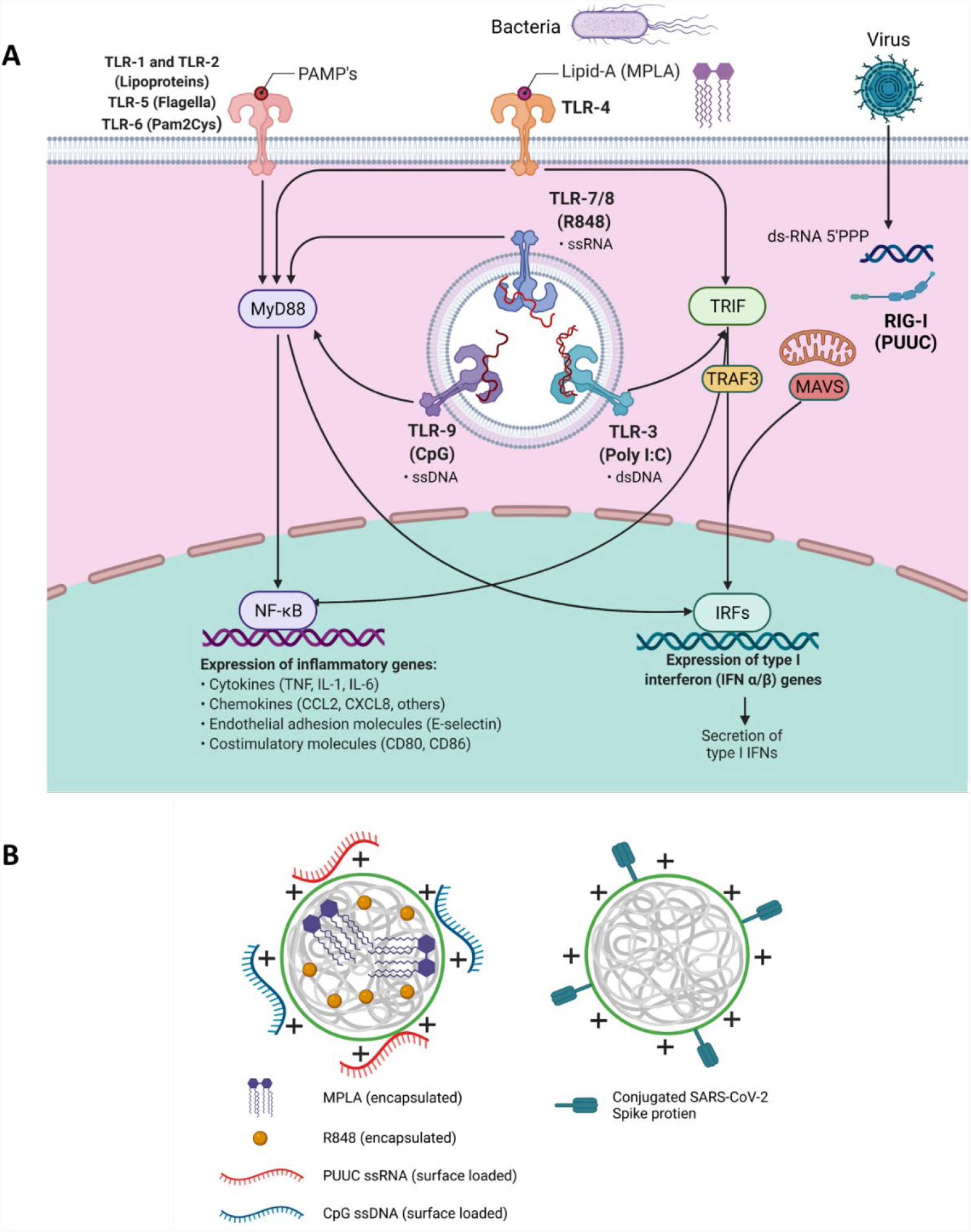
Multiple TLR and RLR intracellular signaling pathways intersect and can be activated by nanoparticles with encapsulated and surface-loaded pathogen-associated molecular patterns (PAMPs). **A)** Schematic of signaling downstream from TLR 1, TLRs 3-9, and RIG-I-like receptors indicating MyD88, TRIF, TRAF3, and MAVS as important intermediates leading to nuclear translocation of NF-kB or IRFs to synthesize proinflammatory genes. **B)** Depiction of poly(lactic-co-glycolic acid)-polyethyleneimine nanoparticles (PLGA-PEI NPs) with encapsulated hydrophobic molecules (R848, MPLA) and/or surface loaded nucleic acids (CpG, PUUC) for adjuvant delivery or surface-loaded SARS-CoV-2 spike protein for antigen delivery.

A single pathogen can express multiple PAMPs which concurrently stimulate multiple PRRs on immune cells. For instance, the live-attenuated yellow fever vaccine activates TLR2, TLR7, TLR8, and TLR9 on different DC subsets. Similarly, the live-attenuated Bacillus Calmette–Guérin (BCG) tuberculosis vaccine signals through TLR2, 4, and 9, among other PRRs.^20^ Positive-sense single-stranded RNA viruses like SARS-CoV-2 interact with TLR7 and TLR8, and produce dsRNA replication intermediates that are recognized by RIG-I and TLR3.^21–24^ TLR2 (dimerized with TLR1 or TLR6) and TLR4 have also exhibited potential to recognize viral proteins; in fact, SARS-CoV-2 S protein reportedly induces TLR4 signaling.^25–27^ Studies have shown that combinations of PAMPs result in synergistic, complementary, and antagonistic effects on innate and adaptive immunity. We have recently reported co-delivery of MPLA and CpG DNA on poly(lactic-*co*-glycolic acid)-polyethylenimine (PLGA-PEI) particles induces prolonged IRF-5 phosphorylation, leading to synergistic increases in proinflammatory cytokine secretion in APCs.^28^ Further, we have demonstrated that intramuscular vaccination of R848, PUUC and H1N1 hemagglutinin on PLGA-PEI particles elevates CD8+ T cell populations in the lung and CD4+ antigen-specific T cell populations in the spleen in mice.^29^

To evaluate the effects of PRR-targeted adjuvants on SARS-CoV-2 protein subunit vaccination, we first formulated PLGA-PEI nanoparticles (NPs) with pairings of MPLA (TLR4), R848 (TLR7/8), or CpG DNA (TLR9) plus PUUC (RIG-I), and then combined the NPs with SARS-CoV-2 spike S1 subunit (containing the ACE-2 receptor binding domain) for in vitro studies or full stabilized S protein for in vivo studies. The various formulations were tested in vitro using iso-mixed leukocyte reactions (iso-MLRs) and in vivo using intranasal and intramuscular prime-boost vaccination models. SARS-CoV-2 is an airborne virus that is mainly transmitted through contact with respiratory droplets from the nose and throat of an infected person^30,31^, resulting in severe lung infection and subsequent systemic infection (i.e., viremia)^32^. We explored both intranasal vaccination (I.N.) to induce local immune memory in the lung that could protect from SARS-CoV-2 infection and viremia as well as prevent transmission, and intramuscular (I.M.) vaccination to induce broad systemic immunity. We show that I.N. vaccination with MPLA+PUUC NP induced CD4+CD44+ activated memory T cells in the lung while MPLA NPs produced anti-S protein IgG and IgA antibodies in the broncho-alveolar lavage (BAL) fluid. Interestingly, I.N. vaccination did not produce detectable antibodies in the serum. In contrast PUUC-carrying NPs, when delivered I.M. with various doses of the S protein, produced strong humoral responses, characterized by increased neutralizing antibody levels and germinal center (GC) B cells. Intramuscular vaccination of MPLA+PUUC NP generated similar responses to PUUC NP while further raising anti-spike protein neutralization titers and increasing T follicular helper cell responses in the draining lymph nodes (dLNs). Collectively, our data show route-specific polarization of local versus systemic immune responses against SARS-Cov2 using NP-adjuvanted protein subunit vaccines and suggest potential for inducing both lung-specific and systemic immunity by targeting I.N. vs I.M vaccination routes.

## MATERIALS AND METHODS

### Synthesis of PUUC (RIG-I Agonist)

The RIG-I agonist poly-U/UC (PUUC) is based on a Hepatitis C Virus (HCV) RNA sequence.^10^ PUUC RNA was transcribed from custom DNA templates (Integrated DNA Technologies, Custom PAGE-purified Ultramer oligos) with the MEGAshortscript T7 Transcription Kit (Invitrogen, Cat# AM1354) as previously described.^19^ The templates were: 5’-TAA TAC GAC TCA CTA TAG GCC ATC CTG TTT TTT TCC CTT TTT TTT TTT CTT TTT TTT TTT TTT TTT TTT TTT TTT TTT TTT TTC TCC TTT TTT TTT CCT CTT TTT TTC CTT TTC TTT CCT TT-3’ (forward) and 5’-AAA GGA AAG AAA AGG AAA AAA AGA GGA AAA AAA AAG GAG AAA AAA AAA AAA AAA AAA AAA AAA AAA AAA AAA AGA AAA AAA AAA AGG GAA AAA AAC AGG ATG GCC TAT AGT GAG TCG TAT TA-3’ (reverse). PUUC was purified by chilled ethanol precipitation followed by centrifugation and resuspension in nuclease free water (Boston BioProducts, Cat# R-100DR) at 0.5 to 1.0 mg/mL, and yields were quantified the Nucleic Acid Quantification workflow on a Synergy HT plate reader (BioTek) with Gen5 software. Synthesized PUUC was stored at −80°C until use.

### PLGA-PEI Nanoparticle Synthesis and Adjuvant Loading

PLGA nanoparticles were synthesized via a double-emulsion and solvent evaporation method as previously reported.^28,29,33^ PLGA (50:50, MW: 7,000-17,000, Resomer RG 502H, Sigma-Aldrich, Cat# 719897) was dissolved in dichloromethane (DCM, Sigma-Aldrich, Cat# 270997) in 1:20 w/v ratio. Endotoxin-free water was added in a 1:4 v/v ratio to the mixture to form the first water-oil emulsion. For particles with encapsulated adjuvant, either R848 (4.80-6.67 μg/mg PLGA, STEMCELL Technologies, Cat# 73784) or MPLA PHAD® (1 μg/mg PLGA, Avanti Polar Lipids, Cat# 699800P) was dissolved the DCM added to the first water-oil emulsion. The first emulsion was sonicated for 2 min at 65% power at RT, and then added to 5% PVA (MW: 31,000-50,000) in water at a 5:16 v/v ratio to form a second water-oil-water emulsion. The second emulsion was sonicated at 65% power for 5 min at RT. DCM was evaporated by stirring the second emulsion at RT for 3 h. Large PLGA aggregates were removed by centrifugation at 2000 x g for 10 min. The supernatant was then ultracentrifuged at 80,000 x g for 20 min to pellet the PLGA NPs. Nanoparticles were washed with DI water via ultracentrifugation, resuspended in DI water and lyophilized for 48 h. Branched polyethyleneimine (bPEI, Polysciences, Cat# 06090) was coated onto the PLGA nanoparticles by EDC (Thermo Scientific, Cat# 22980) and sulfo-NHS (Thermo Scientific, Cat# PG82071) chemistry to produce cationic PLGA-PEI nanoparticles. PLGA NPs were resuspended in 0.1 M aqueous MES (Sigma-Aldrich, Cat# M5287) and 40 molar excess of EDC and 25 molar excess of sulfo-NHS were added to the suspension. After end-to-end rotation for 2 h, a 1:2 v/v bPEI solution in 0.2 MES was added to the particle suspension. After stirring for 2 h, particles were ultracentrifuged at 80,000 x g twice in 1 M NaCl and once in DI water. PLGA-PEI NPs were resuspended in DI water and lyophilized for 48 h. For particles electrostatically loaded with nucleic acid adjuvants, either Class B CpG ODN 1826 (Invitrogen, Cat# tlrl-1826) or PUUC was incubated with particles in 10 mM sodium phosphate buffer (made with nuclease free water) under end-to-end rotation for 24 h.

Nanoparticle size and surface zeta potential were measured with a Zetasizer Nano ZS (Malvern). R848 loading was determined by dissolving particles in sterile-filtered DMSO (Tocris, Cat# 3176) followed by absorbance readings against a standard curve at 324 nm. MPLA loading estimation via GC-MS, LC-MS, and surrogate fluorometry has been previously described.^28^ PUUC and CpG loading was quantified by supernatant measurement of unbound RNA or DNA, respectively, with the Nucleic Acid Quantification workflow on a Synergy HT plate reader (BioTek) with Gen5 software.

### Spike-Conjugated Nanoparticle Synthesis

To conjugate SARS-CoV-2 spike to PLGA-PEI NPs, free amines of particles were converted to thiols via Traut’s reagent. Initially, PLGA-PEI NPs were dispersed in 100 mM phosphate buffer with EDTA (2 mM) (pH = 8.0) and mixed with excess amount of 2-iminothiolane (Traut’s reagent, Sigma-Aldrich, Cat# I6256). After continuous rotation overnight at RT, thiolated PLGA-PEI (PLGA-PEI-SH) NPs were purified by centrifugation, washing and lyophilization overnight. Thiol groups on particles were estimated by Ellman’s reagent (G Biosciences, Cat# BC87) using cysteine (Sigma-Aldrich, Cat# 168149) as the standard, yielding a 60-70% thiolation efficiency. Until spike loading, PLGA-PEI-SH NPs were stored at −20°C. Prior to *in vivo* experiments, a stock solution of NHS-PEG-SPDP (bifunctional crosslinker, Sigma-Aldrich Cat# 803499) was prepared in anhydrous DMSO (10 mg/mL, Sigma-Aldrich, Cat# 276855). Further, excess NHS-PEG-SPDP (200 μg) crosslinker was added to well-dispersed PLGA-PEI-SH NPs solution in PBS (5 mg/mL, pH = 7.6) with stabilized SARS-CoV-2 spike glycoprotein (10 μg/mg NP, with Avi Tag, BEI Resources, Cat# NR-53524). Following rotation for 6 h at RT, NPs were centrifuged at 21,000 x *g* for 10 min, and supernatant was saved (first wash). NPs were washed with nuclease free water, centrifuged, and supernatant was also saved (second wash). Supernatant washes were filtered through a 100,000 MWCO Ultra-4 Amicon centrifugal filter (Millipore Sigma, Cat# UFC810096) to retain unconjugated spike glycoprotein and separate remaining NHS-PEG-SPDP crosslinker. Unconjugated spike protein was measured via micro-BCA (Boster Bio, Cat# AR1110).

### Bone Marrow Derived Dendritic Cell (BMDC) Culture

GM-CSF-derived BMDCs were generated similarly to other reports.^34^ Female Balb/cJ (5-10 weeks old, Jackson Labs) were euthanized with CO^2^. PBS was flushed through tibiae and femurs to isolate bone marrow, which was strained through a 40 μm strainer. Bone marrow cells were centrifuged, treated in RBC lysis buffer (Thermo Fisher, Cat# 00-4333-57), and were seeded in 100-mm round tissue-culture treated polystyrene dishes at 1 million cells/mL in complete RPMI medium (with L-glutamine, Gibco, Cat# 11875119) with 10% characterized fetal bovine serum (GE Healthcare, Cat# SH30071.03), 1% penicillin-streptomycin (Corning, Cat# 30-002-CI), 1xβ-mercaptoethanol (Thermo Fisher, Cat# 21985023), 1 mM sodium pyruvate (Thermo Fisher, Cat# 11360070), and 20 ng/mL murine GM-CSF (Peprotech, Cat# 315-03). Media was replenished on days 2, 4, and 6 of culture, and experiments with GM-CSF BMDCs were conducted on day 7 of culture. For FLT3L BMDCs, RBC-lysed murine bone marrow was seeded at 2 million cells/mL in complete RPMI media with 200 ng/mL human FLT3 ligand (PeproTech, Cat# 300-19) in a 6-well plate (10 million cells/well). Experiments with FLT3L BMDCs were conducted on day 9 of culture.

### Characterization of APC Subsets in BMDC Culture

Both FLT3L and GM-CSF BMDCs were stained with 1:1000 Zombie Green Fixable Viability Kit (BioLegend, Cat# 423111) and were blocked with anti-mouse CD16/32 (BioLegend, Cat# 101320) and True-Stain Monocyte Blocker (BioLegend, Cat# 426102). After blocking, cells were stained with anti-mouse CD11b (BUV395, BD, Cat# 563553), CD11c (Brilliant Violet 421, Biolegend, Cat# 117343), B220 (APC, Biolegend, Cat# 103212), Ly-6C (Brilliant Violet 711, Biolegend, Cat# 128037), Ly-6G (Brilliant Violet 785, Biolegend, Cat# 127645), CD64 (PE, Biolegend, Cat# 139304), F4/80 (PE/Cy5, Biolegend, Cat# 123112), and MHC-II (APC/Cy7, Biolegend, Cat# 107628). Cells were fixed with BD Cytofix (Cat# 554655) and analyzed with a BD LSRFortessa flow cytometer. FlowJo Software (BD) was used to generate t-distributed stochastic neighbor embedding (t-SNE) plots.

### In Vitro Activation of BMDCs and Iso-Mixed Lymphocyte Reaction (Iso-MLR)

Adjuvanted PLGA-PEI NPs and the S1 subunit of the SARS-CoV-2 spike protein (Novus Biologicals, Cat# NBP2-90985, 100 ng protein/500,000 cells/mL RPMI media) were added to GM-CSF or FLT3L-cultured BMDCs in a U-bottom 96 well plate. Each well contained 100,000 BMDCs in 200 mL media. Doses of adjuvants are outlined in **Table 1**. The concentration of particles was adjusted based on the adjuvant dose and the particle-only (“blank”) control was matched to the highest PLGA-PEI NP concentration to ensure NPs were not activating BMDCs. The cells were incubated with adjuvant-NPs for 24 hours before addition of T cells. To isolate T cells, spleens from female Balb/cJ mice were dissociated in 2 mg/mL Collagenase D (Sigma-Aldrich, Cat# 11088882001) in Opti-MEM media (Gibco, Cat# 11058021) and filtered through a 40 μm cell strainer. T cells were magnetically separated from other splenocytes using the Mouse Pan-T Cell Isolation Kit II (Miltenyi Biotec, Cat# 130-095-130), labeled with CellTrace™ CFSE (Thermo Fisher, Cat# C34554), and resuspended in complete RPMI medium at 1 million cells/mL. BMDC-PLGA NP U-bottom plates were centrifuged at 500 x g for 5 minutes and supernatants were collected and frozen at −20°C. 200,000 T cells were added to each well containing BMDCs. After 72 hours, T cells were stained for flow cytometric analysis with the following: Zombie UV (Biolegend, Cat# 423107) to exclude dead cells, and stained with anti-mouse CD3 (Brilliant Violet 786, Biolegend), anti-mouse CD4 (APC, Biolegend) and anti-mouse CD8a (PE, Biolegend) to identify CD4+ and CD8+ T cells. A well containing only CSFE-stained T cells was included as a non-proliferative control. Percentages of proliferating T cells were calculated by gating on the reduced FITC signal (Figure S1). DuoSet ELISA kits (R&D Systems) were used to quantify IFN-β and IFN-λ3 from BMDC supernatants. IL-12p70, IL-1β, and IL-27 were quantified using Luminex® assays (R&D systems). Supernatants were diluted 4-fold for all cytokines except for IL-1β, in which case they were diluted 8-fold. Supernatants were analyzed in triplicate for each experimental condition.

**Table 1.** Single and combination adjuvant loading on PLGA NP and doses for GM-CSF and FLT3L BMDCs in iso-MLR assays.

| Particle Adjuvants | Adjuvant combinations | Loading level (µg/mg PLGA-PEI NP) | Adjuvant Dose for BMDCs (ng/500,000 cells/mL) |
| --- | --- | --- | --- |
| PLGA-CpG | CpG | 60 | 500 |
| PLGA-MPLA | MPLA | 6 | 100 |
| PLGA-PUUC | PUUC | 6 | 100 |
| PLGA-R848 | R848 | 4.8 | 100 |
| PLGA-CpG+PUUC | CpG | 30 | 500 |
|  | PUUC | 6 | 100 |
| PLGA-MPLA+CpG | MPLA | 6 | 50 |
|  | CpG | 60 | 500 |
| PLGA-MPLA+PUUC | MPLA | 6 | 100 |
|  | PUUC | 6 | 100 |
| PLGA-R848+PUUC | R848 | 4.8 | 100 |
|  | PUUC | 5 | 100 |

### In Vivo Intranasal Vaccination

For this experiment, there were six mice per treatment and control group for a total of 78 mice in thirteen groups: **(A)** Seven vaccine groups included 1 μg of unformulated SARS-CoV-2 spike protein (R&D Systems, Cat# 10549-CV-100) delivered with adjuvanted NPs: MPLA (24 μg), CpG (20 μg), PUUC (17 μg), CpG-PUUC (20 μg, 17 μg), MPLA-PUUC (20 μg, 17 μg), R848-PUUC (20 μg, 20 μg), and MPLA-CpG (24 μg, 20 μg). **(B)** Two vaccine groups included 1 μg of PLGA NP-conjugated SARS-CoV-2 spike protein (methods described above) delivered with adjuvanted NPs: PUUC (17 μg) and MPLA-PUUC (20 μg, 17 μg). **(C)** There were three controls included: saline, unformulated SARS-CoV-2 spike only (1 μg) and PLGA NP-conjugated SARS-CoV-2 spike only (1 μg). **(D)** One vaccine group included 2 μg of unformulated stabilized SARS-CoV-2 spike protein delivered with MPLA-PUUC PLPs (20 μg, 17 μg). On days 0 (prime) and 28 (boost), adjuvant-loaded NPs and either unformulated spike protein or spike-conjugated NPs were resuspended in normal saline and administered to 9–10-week-old female BALB/c mice dropwise in the bilateral nares (4 mg NPs and 1 μg antigen in 60 μL saline per mouse). Half of each experimental group (3 mice) was euthanized at day 27, and blood and bronchoalveolar lavage (BAL) fluid was collected. The remaining 3 mice in each group were euthanized at day 35, and blood, BAL fluid, and lungs were collected. All blood samples were clotted for 30-60 min at RT in serum separator tubes (BD, Cat# 365967), and sera were isolated by centrifugation at 4,000 x *g* for 15 min at 4 °C. Sera were heat inactivated at 56 °C for 30 min in a water bath to inhibit complement binding and then stored at −80 °C until use. BAL fluid samples were centrifuged to remove cells and stored at −80°C.

### Ex Vivo Lung Cell Restimulation

Harvested lungs from intranasally vaccinated mice were processed into single cell suspensions with a gentleMACS™ Octo Dissociator and Lung Dissociation Kit (Miltenyi Biotec) according to manufacturer’s instructions including RBC lysis. Cells were centrifuged and resuspended at 10 million cells/mL in RPMI media with 10% FBS, 1% penicillin-streptomycin, 1 mM sodium pyruvate, and 1x β-mercaptoethanol. Cells were seeded at 2.4 million cells per well in a treated 96-well plate and left to culture overnight. Lung cells were centrifuged and resuspended with fresh media with 20 μL/mL of PeptTivator® SARS-CoV-2 Prot_S (Miltenyi Biotec) and 5 μg/mL Brefeldin A (BioLegend). After 6 h, cells were stained for 30 min at RT with Zombie Green™ Fixable Viability Kit (BioLegend) and were blocked with anti-mouse CD16/32 (BioLegend) and True-Stain Monocyte Blocker™ (BioLegend). Following blocking, cell surfaces were stained for 30 min at 4°C with anti-mouse CD45 (BD, BUV395), CD44 (Biolegend, BV711), CD69 (Biolegend, BV785), CD103 (Biolegend, PE-Dazzle 594), CD8a (Biolegend, PE/Cy5), and CD4 (Biolegend APC). Surface-stained cells were fixed and permeabilized for 30 min at 4°C with the Foxp3/Transcription Factor Staining Buffer Set (eBioscience). Then cells were stained with anti-mouse IFN-γ (PE), Granzyme B (Pacific Blue), and TNF-α (PE/Cy7). Staining was measured with a BD LSRFortessa™ flow cytometer.

### In Vivo Intramuscular Vaccination and Analysis of Popliteal Lymph Nodes

NPs were loaded with MPLA (6 μg/mg NP), PUUC (5 μg/mg NP), or MPLA+PUUC. Female BALB/c mice were intramuscularly injected with adjuvant-loaded NPs (4 mg/mouse) and variable doses of stabilized SARS-CoV-2 spike glycoprotein (BEI Resources, Cat# NR-52397) into the left and right anterior tibialis anterior muscles (50 μL NPs in saline per injection) on day 0 (prime) and day 28 (boost). On day 36, mice were euthanized, and blood, bilateral popliteal lymph nodes, and spleens were harvested. Blood was processed as described above to isolate sera. Lymph nodes from each leg per mouse were combined and passed through a 40 μm cell strainer to generate single cell suspensions which were centrifuged and washed with PBS. Lymph node cells were stained with Zombie Green™ (Biolegend) and blocked with anti-mouse CD16/32 as above. After blocking, cell surfaces were stained for 30 min at 4°C with anti-mouse B220 (Biolegend, APC), GL7 (Biolegend, PE/Cy7), and CXCR5 (Biolegend, BV421). Surface-stained lymph node cells were then fixed, permeabilized and intracellularly stained with anti-mouse BCL6 (Biolegend, PE) in permeabilization buffer. Lymph node cells were fixed and analyzed with a BD FACSymphony™ A5 Cell Analyzer. The same experiment was repeated using spike-conjugated nanoparticles instead of unformulated spike protein with 1000 ng spike protein/mouse delivered on days 0 and 28 for prime-boost regimen.

### Ex Vivo Splenocyte Restimulation

Harvested spleens from intramuscularly vaccinated mice were processed into single cell suspensions with a gentleMACS™ Octo Dissociator and Spleen Dissociation Kit (Miltenyi Biotec) with slight modifications to the manufacturer’s protocol. Specifically, to protect splenocyte viability, we ensured that tissue processing began within approximately 30 min of harvesting. As soon as samples were counted, samples were immediately centrifuged and resuspended at 10 million cells/mL in BMDC media. Cells were seeded at 2 million cells per well in a treated 96-well plate and left to culture overnight at 37°C in 5% CO^2^. Cells were centrifuged and resuspended with fresh media with 40 μL/mL of PeptTivator® SARS-CoV-2 Prot_S (Miltenyi Biotec, Cat# 130-126-700), 1 μL/mL monensin (BioLegend), and 5 μg/mL Brefeldin A (BioLegend). After 6 h of incubation, splenocytes were stained for 30 min at RT with Zombie Green™ Fixable Viability Kit (BioLegend) and were blocked with anti-mouse CD16/32 (BioLegend). Following blocking, cell surfaces were stained for 30 min at 4°C with anti-mouse CD8a (Biolegend, PE/Cy5), CD4 (Biolegend, APC), CD3 (BD, BUV395), CD44 (Biolegend, BV711). Surface-stained cells were fixed and permeabilized for 30 min at 4°C with the Foxp3/Transcription Factor Staining Buffer Set (eBioscience). Then cells were stained with anti-mouse IFN-γ (Biolegend, PE), Granzyme B (Biolegend, BV421), IL-4 (Biolegend, PE/Dazzle 594) and TNF-α (Biolegend, PE/Cy7). Staining was measured with a BD FACSymphony™ A5 Cell Analyzer.

### ELISA Assay for Quantifying Anti-Spike Antibody Responses

SARS-CoV-2 stabilized spike (BEI Resources, Cat# NR-52397) was diluted 1 μg/mL in 0.05 M carbonate-bicarbonate buffer (pH 9.6). Diluted spike was adsorbed onto Nunc™ MaxiSorp™ ELISA Plates by incubating 100 ng/well overnight at 4°C. Antigen-coated plates were washed three times with wash PBST (0.01 M PBS + 0.05% Tween-20), and plates were blocked for 6-8 h at 4°C with PBSTBA (PBST + 1% BSA + 0.02% NaN^3^). Blocked plates were incubated overnight at 4°C with 10^2^- to 10^6^-fold diluted BAL fluid or sera from intranasally or intramuscularly vaccinated mice, respectively. Plates were washed three times with PBST. A secondary biotinylated anti-mouse IgA, total IgG, IgG1, or IgG2a antibody (SouthernBiotech) was diluted 5,000-fold in 5-fold diluted PBSTBA and was added to plates for 2 h at RT. Plates were similarly washed with PBST. 5,000-fold diluted streptavidin-conjugated horseradish peroxidase (strep-HRP, ThermoFisher) was incubated for 2 h at RT. Plates were washed six times. Ultra TMB-ELISA Substrate Solution (ThermoFisher) was incubated for 15 to 25 min for color to develop on the plate. Lastly, 2 N sulfuric acid was added to each well, and absorbance was measured at 450 and 630 nm on a Synergy HT plate reader (BioTek) with Gen5 software.

### Modified ELISA Assay to Measure Anti-Spike Neutralizing Antibodies

Neutralizing antibodies were quantified like the ELISA method described above in a 384-well UltraCruz® ELISA high-binding plate (Santa Cruz Biotechnology). Diluted spike in 0.05 M carbonate-bicarbonate buffer (1 μg spike/mL, pH 9.6) was incubated in wells of a 384-well plate (50 ng/well) overnight at 4°C. Plates were washed three times with PBST, and plates were blocked in PBSTBA for 6-8 h at 4°C. Blocked plates were incubated overnight at 4°C with 10e2- to 10e6-fold diluted sera from intramuscularly vaccinated mice. Plates were similarly washed. In lieu of a secondary antibody, plates were incubated for 2 h at RT with 500 ng/mL (25 ng/well) recombinant biotinylated human ACE-2 (R&D Systems, Cat# BT933-020) diluted in PBSTBA. Plates were washed with PBST, and 5,000-fold diluted strep-HRP (ThermoFisher) was incubated for 2 h at RT. Plates were washed again six times, and incubated with Ultra TMB-ELISA Substrate Solution (ThermoFisher) for 20 min. The reaction was stopped with 2 N sulfuric acid, and absorbance was measured at 450 and 630 nm on a Synergy HT plate reader (BioTek) with Gen5 software. Absorbances were normalized by row to correct for plate-based effects.

### Statistical Analysis

All flow cytometry FCS files were analyzed with FlowJo (v10, BD). Statistical analyses were performed with GraphPad Prism 9. Normality was assessed with the Kolmogorov-Smirnov test with Dallal-Wilkinson-Lilliefors *P* value. For more than two comparisons, statistical differences between normally distributed datasets were determined with a one-way ANOVA with Tukey’s post-hoc test for multiple comparisons. Similarly, nonparametric datasets were evaluated with the Kruskal-Wallis test and Dunn’s post-hoc test. For antibody quantification with ELISA, area under the curve (AUC) across fold dilutions was computed with the AUC function on GraphPad as previously reported.^35^

## RESULTS

### TLR- and RIG-I-targeted combination adjuvants differentially induce GM-CSF BMDC proinflammatory cytokine secretion and FLT3L BMDC activation of T cells when delivered with SARS-CoV-2 S1 subunit protein

Hydrophobic R848 or MPLA were encapsulated into PLGA NP using a w/o/w emulsion-solvent evaporation method previously published. Blank particles without adjuvants were also prepared as a negative control. The average diameter of PLGA NPs prior to PEI modification was approximately 250 nm.^28,29,33^ To electrostatically load CpG DNA and PUUC RNA, PLGA NPs were modified with surface bPEI to produce cationic particles with an approximate zeta potential of 30 mV.^28,29,33^ To assess if adjuvants are more immunostimulatory when paired with another or delivered alone, combination adjuvants MPLA+PUUC, CpG+PUUC, and R848+PUUC on NPs as well as single adjuvant and unformulated (Blank) control NPs were mixed with SARS-CoV-2 S1 subunit in media and incubated (adjuvant and S1 doses in Experimental Methods section) with murine BMDCs generated using either GM-CSF or FLT3L cytokines in a 96 well plate. After 24 hours, supernatants were collected to quantify BMDC proinflammatory cytokine secretion, and then T cells were added for another 72 hours with activated BMDCs before quantifying T cell proliferation using CellTrace CFSE.

Flow cytometry and tSNE analysis revealed Day 9 FLT3L-derived BMDC cultures were composed of conventional dendritic cells (cDC; CD11c^+^MHCII^hi^CD64^lo/–^F4/80^lo/–^) and plasmacytoid dendritic cells (pDC; B220^+^Ly6C^+^CD11c^+^MHCII^lo^Ly6G^lo^), whereas Day 7 GM-CSF-derived BMDCs were primarily monocytes (Ly6C^hi^CD11c^−^) and monocyte-derived DCs/macrophages (MoDC; CD64^+^F4/80^+^). There is some macrophage survival in FLT3L culture, and neutrophil (Ly6G^hi^Ly6C^+^) survival in GM-CSF culture (**Figure 2A**). MPLA+PUUC NPs statistically increased IL-1β and IL-27 secretion, R848+PUUC NPs upregulated IL-12p70 and IL-27, and CpG+PUUC NPs increased production of IFN-β in GM-CSF BMDC culture compared to single adjuvant NP, blank NP, and antigen-only controls (**Figure 2B-F**). In FLT3L BMDC culture, however, CpG NPs increased and CpG+PUUC NPs antagonistically decreased IFN-λ3 secretion (**Figure 2G**).

**Figure 2:**
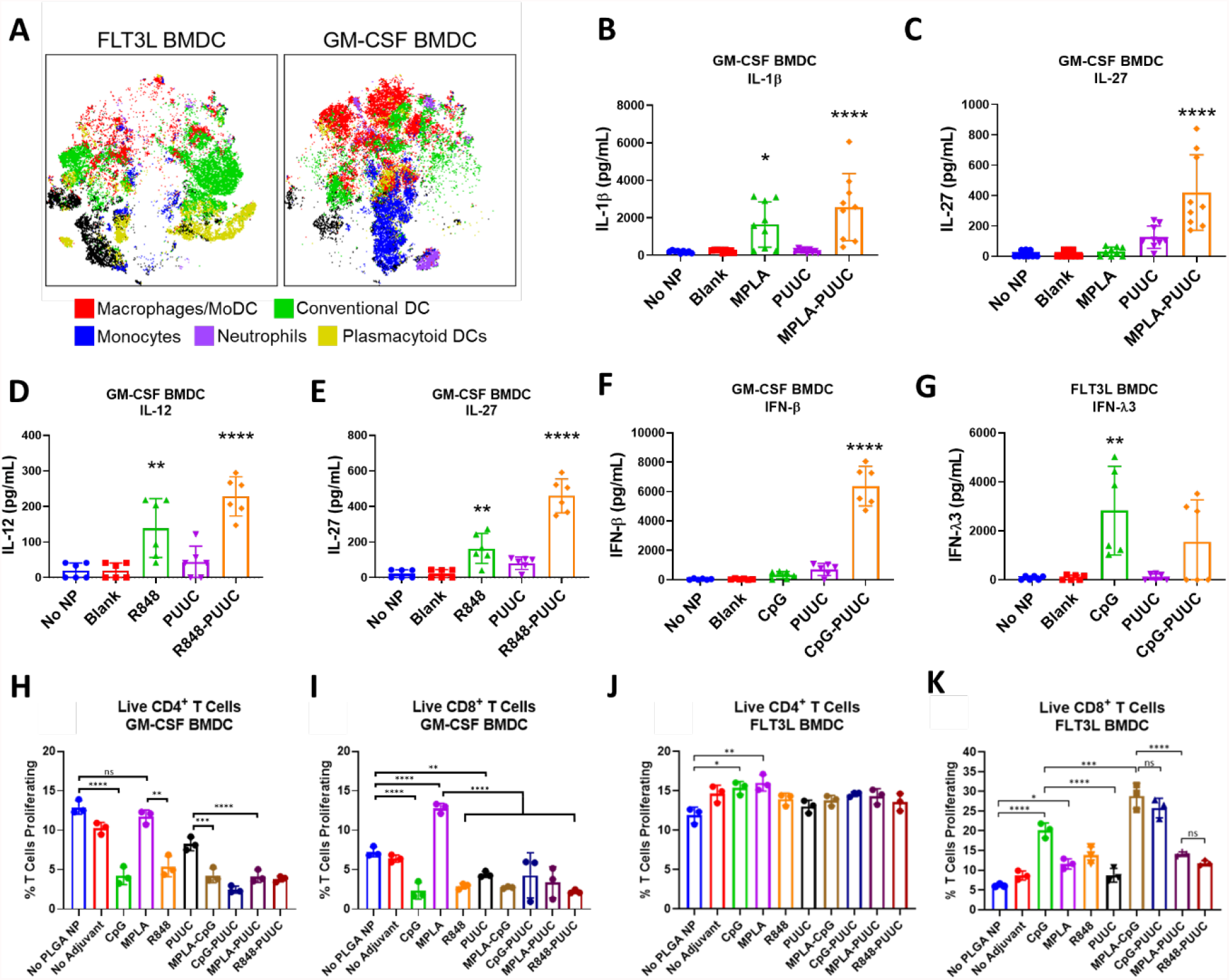
GM-CSF and FLT3L BMDCs secrete different cytokine profiles and activate CD8+ T cells in response to S1 protein with combination adjuvants. **A)** t-SNE plots of GM-CSF and FLT3L BMDCs with labeled clusters of APC subsets. Macrophages/Mo-DCs are CD64^+^ F4/80^-^, Conventional DCs are CD11c^+^ MHCII^+^ CD64^lo^ F4/80^lo^, Monocytes are Ly6C^hi^ CD11c^-^, Neutrophils are Ly6G^hi^ Ly6C^+^, plasmacytoid DCs are B220^+^ Ly6C^+^ CD11c^+^ MHCII^lo^. **B-G)** Cytokine concentrations (pg/mL) of IL-1β, IL-27, IL-12p70, IFN-β, and IFN-λ3 in supernatants of BMDC culture after incubation with adjuvanted NPs for 24 h. **H-I)** Percentage of live CD3+CD4+ T cells or CD3+CD8+ T cells proliferating in presence of GM-CSF BMDCs, gated on diminished CFSE signal (Figure S1). “No Adjuvant” condition is non-adjuvant (blank) NPs. **J-K)** Percentage of live CD3+CD4+ T cells or CD3+CD8+ T cells proliferating in presence of FLT3L BMDCs. *p<0.05. **p<0.01, ***p<0.001, ****p<0.0001 based on a One-Way ANOVA and Tukey Test for multiple comparison.

Next, we evaluated the ability of combination adjuvant-activated BMDCs to stimulate murine T cell proliferation in iso-mixed lymphocyte (iso-MLR) reactions. After 72 h there were no significant changes in CD4 T cell proliferation with MPLA- or PUUC-stimulated GM-CSF BMDCs while CpG, R848, MPLA-CpG, CpG+PUUC, MPLA+PUUC and R848+PUUC-stimulated GM-CSF BMDCs induced two-fold decreases in proliferation compared to antigen-only controls (**Figure 2H**). Alternatively, there was a two-fold increase in CD8 T cell proliferation with MPLA-stimulated GM-CSF BMDCs (**Figure 2I**). CD4 T cell proliferation significantly increased when mixed with CpG NP- and MPLA NP-stimulated FLT3L BMDCs (**Figure 2J**). CD8 T cell proliferation significantly increased when mixed with FLT3L BMDCs stimulated with CpG, MPLA-CpG, and CpG+PUUC NPs (**Figure 2K**). Combination adjuvants synergistically or additively induced proinflammatory cytokine secretion by APCs in vitro while select adjuvants, including MPLA NP with GM-CSF BMDCs and CpG-single and dual NP formulations with FLT3L BMDCs, increased CD8 T cell proliferation *in vitro*. We were motivated to evaluate whether these *in vitro* studies would predict how combination and single TLR- and RIG-I targeted adjuvants would improve SARS-CoV-2 subunit vaccination *in vivo*.

### MPLA-PUUC NPs increase lung T cell responses in mice following intranasal vaccination with SARS-CoV-2 stabilized spike protein

Given the protective role of mucosal immunity against respiratory virus SARS-CoV-2,^36^ we vaccinated mice intranasally with single and dual adjuvant-loaded NPs combined with either soluble (S-S) or NP-conjugated S protein (NP-S). Mice were intranasally immunized with a primer on Day 0 and booster of the same dose on Day 28. Lungs were harvested and cell lysates restimulated with S-S and NP-S on Day 35. Lungs from mice immunized with MPLA+PUUC NP plus NP-S had significantly higher CD4^+^CD44^+^ T cell populations with significant increases in IFNγ and TNFα secretion when restimulated with S peptide pools (**Figures 3A-C**). CD69^+^CD103^+^ tissue-resident memory T cell populations were highest in mice vaccinated with MPLA NP and MPLA+PUUC NP plus S-S (**Figure 3D**). These cells exhibited a double negative CD4^−^CD8^−^ phenotype, suggesting they could be γδ cells, a subset of T cells enriched in epithelial and mucosal tissues that are activated in an MHC-independent manner. Intranasal vaccination with CpG, CpG-PUUC, R848-PUUC, or MPLA-CpG NP delivered with S-S failed to generate significant increases in percentages of T cells producing IFNγ or TNFα (**Figure S2A-K**). In mice immunized with MPLA NP plus S-S, anti-S BAL fluid IgG and IgA and serum IgG levels were higher compared to mice in PUUC NP and MPLA+PUUC NP groups **(Figures 2E-G)**.

**Figure 3:**
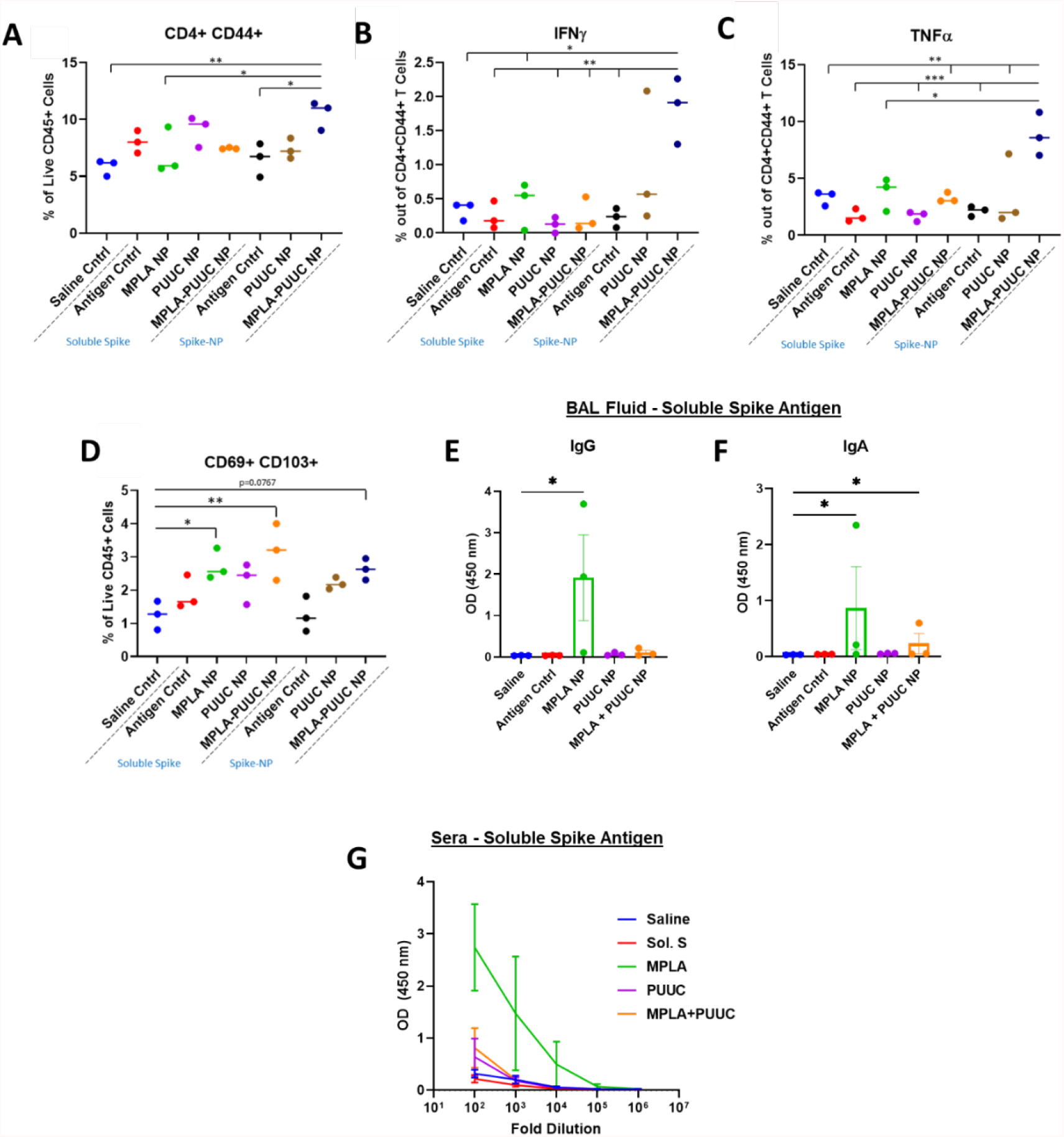
MPLA+PUUC NPs increase T cell responses in the lung when delivered intranasally with spike protein. On days 0 (prime) and 28 (boost), female BALB/c mice were immunized by dropwise addition of saline and spike protein (1 μg) with PLGA-PEI NPs (4 mg) loaded with MPLA (24 μg), PUUC (17 μg), and MPLA+PUUC (20 μg, 17 μg). The same adjuvant-NPs were combined with spike-conjugated NPs. Mice were euthanized and lungs were collected on day 35, one week after the booster. Lung cells were restimulated with spike peptide pools for 6 h and stained for analysis by flow cytometry. Percentages of cells expressing **A)** CD4+CD44+ out of CD45+ cells, **B)** IFNγ+ out of CD4+CD44+ cells, **C)** TNFα+ out of CD4+CD44+ cells, **D)** CD69+CD103+ (tissue resident memory T cells) out of CD45+ cells. BAL fluid from vaccinated mice using soluble spike antigen was assayed for anti-spike **E)** IgG and **F)** IgA with ELISA. **G)** Sera were assayed for anti-spike IgG with ELISA (error bars represent the SEM). P values are *p ≤ 0.05, **p ≤ 0.01, ***p ≤ 0.001 calculated using **A-D)** One-way ANOVA with Tukey post-hoc test or **E-F)** Kruskal-Wallis with Dunn’s post-hoc test for nonparametric data.

### PUUC NPs increase humoral responses in mice following intramuscular vaccination with various doses of SARS-CoV-2 stabilized spike protein

Because authorized COVID vaccines in the US are administered intramuscularly and antigen doses have been shown to impact T cell responses,^37^ we evaluated the ability of PUUC NPs to improve SARS-CoV-2 intramuscular protein subunit vaccines with different doses of S protein. On Days 0 and 28, mice were immunized with S protein at doses of 80, 200, or 1000 ng with or without PUUC NPs. A cohort of mice received a mixed dose of 80 ng on Day 0 and 1000 ng on Day 28. Blood was drawn via the jugular vein of mice on Day 26 and cardiac puncture on Day 36 to quantify pre- and post-booster levels of antigen-specific IgG, which were higher in mice with PUUC NP-adjuvanted vaccines (**Figure 4A-D**). Post-booster anti-S IgG1 and IgG2a titers were simultaneously increased, especially for the 200/200, 1000/1000, and 80/1000 (prime/boost) vaccination groups with PUUC NPs (**Figure S3A-D**). Neutralization of S protein with post-booster mouse sera was significantly detectible up in the 200/200, 1000/1000, and 80/1000 ng prime/boost antigen concentrations adjuvanted with PUUC. Up to a 1000-fold dilution of sera, neutralization of spike was significantly higher (ACE-2 A450 absorbance signal lower) in the 1000/1000 ng S mixed with PUUC NP group (**Figure 4E**).

**Figure 4:**
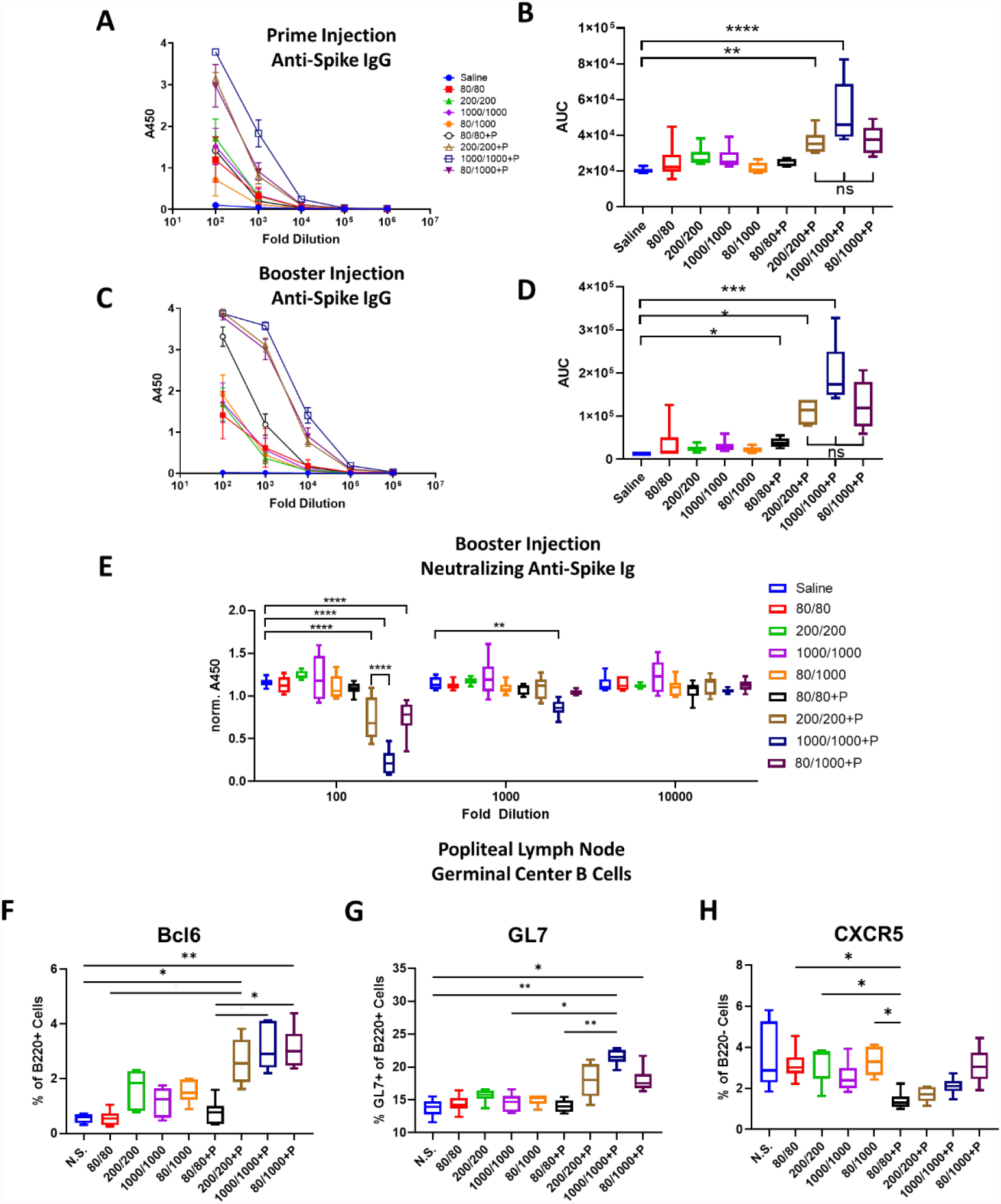
PUUC NPs delivered intramuscularly with spike protein enhance humoral responses. Female BALB/c mice were immunized by intramuscular injection into both tibialis anterior muscles at day 0 (prime) with saline or soluble spike protein at 80 ng, 200 ng, 1000 ng with or without adjuvant-NPs (4 mg) loaded with PUUC (20 ng). Peripheral blood was drawn on day 26. On day 28, mice were injected with the same, except that half of the 80 ng prime cohort were given a 1000 ng booster. Mice were euthanized on day 36 for harvesting blood, splenocytes, and popliteal LNs. **A)** Anti-spike IgG in pre-booster sera at various dilutions measured by absorbance at 450 during ELISA assays and **B)** comparison of area under the curve (AUC). **C-D)** Anti-spike IgG in post-booster sera measured by absorbance at 450 and comparison of AUC. **E)** Neutralizing anti-spike antibody levels, quantified with absorbance at 450 nm in a modified ELISA with biotinylated ACE-2 biotin. For neutralizing antibody quantification, absorbance was normalized to a blank well in each row of a 384 well plate to correct for plate-based effects. Lower absorbance values indicate higher neutralizing antibody levels. Percentages of cells expressing **F)** Bcl6+ out of B220+ cells, **G)** GL7+ out of B220+ cells and **H)** CXCR5+ out of B220-cells from the combined popliteal lymph nodes. **B**,**D)** Normality was assessed with the Kolmogorov-Smirnov test. Statistical significance was determined with the Kruskal-Wallis test and Dunn’s post-hoc test for multiple comparisons. **E-H)** Statistical significance calculated with One-Way ANOVA and Tukey post-hoc test. *p ≤ 0.05, **p ≤ 0.01, ***p ≤ 0.001, ****p ≤ 0.0001 for all graphs.

By Day 36, percentages of GC BCL6^+^B220^+^ cells in popliteal lymph nodes significantly increased in the 200/200, 1000/1000, and 80/1000 ng S protein mice immunized with PUUC NP compared to non-adjuvanted and saline-injected control mice (**Figure 4F**). Similarly, GL7^+^B220^+^ cell populations were significantly increased in mice immunized with 1000/1000 and 80/1000 ng S protein plus PUUC NP compared to controls (**Figure 4G**). PUUC NP with 80/80 ng S protein significantly decreased populations of B220^−^ CXCR5^+^ cells (T follicular helper, Tfh cells) in the dLNs (**Figure 4H**). In the spleen, PUUC NPs did not significantly change CD4 T cell population percentages (**Figure S4A**) nor IFN-γ, IL4, or TNF-α secretion by CD4 T cells (**Figure S4B-D**). CD8+ T cell populations and secretion of Granzyme B and TNF-α also did not change (**Figure S4I-J, L, M-N, P**). Interestingly, CD8+ T cell secretion of IFNγ decreased in mice receiving PUUC NPs (**Figure S4K, O**).

### Combination adjuvant MPLA+PUUC NPs do not increase humoral responses to intramuscular vaccination with SARS-CoV-2 spike protein compared to single adjuvant PUUC NPs

MPLA+PUUC NP increased lung cellular responses after intranasal S protein subunit vaccination and antigen-specific IgG was not significantly different between the 1000/1000 and 80/1000 ng S protein intramuscular PUUC-adjuvanted vaccination groups. Therefore, we evaluated if MPLA+PUUC NP mixed with 80/1000 ng S-protein would enhance immune responses compared to single adjuvants. Mice were immunized with S protein mixed with MPLA NPs, PUUC NPs, and MPLA+PUUC NPs on Days 0 and 28. Pre- and post-booster antigen-specific IgG levels most significantly increased in mice vaccinated with 80 ng S protein plus PUUC NP (**Figure 5A-D**). Neutralization of S protein was detectable with post-booster sera of PUUC NP-vaccinated mice up to a 100-fold dilution and MPLA+PUUC NP-vaccinated mice up to a 10,000-fold dilution (**Figure 5E**). In the draining popliteal LNs, B220+ cell percentage increased approximately 1.5x in the MPLA NP group (**Figure 5F**). Out of B220^+^ cells, BCL6^+^ and GL7^+^ percentages significantly increased in the MPLA, PUUC, and MPLA+PUUC NP groups relative to the saline control group, and in the PUUC NP group relative to the antigen-only control group (**Figure 5G-H**). Out of B220^−^ cells, the CXCR5^+^ population significantly increased in the PUUC and MPLA+PUUC NP groups compared to the saline control and in the MPLA+PUUC NP group relative to the antigen-only control group **(Figure 5I**).

**Figure 5.**
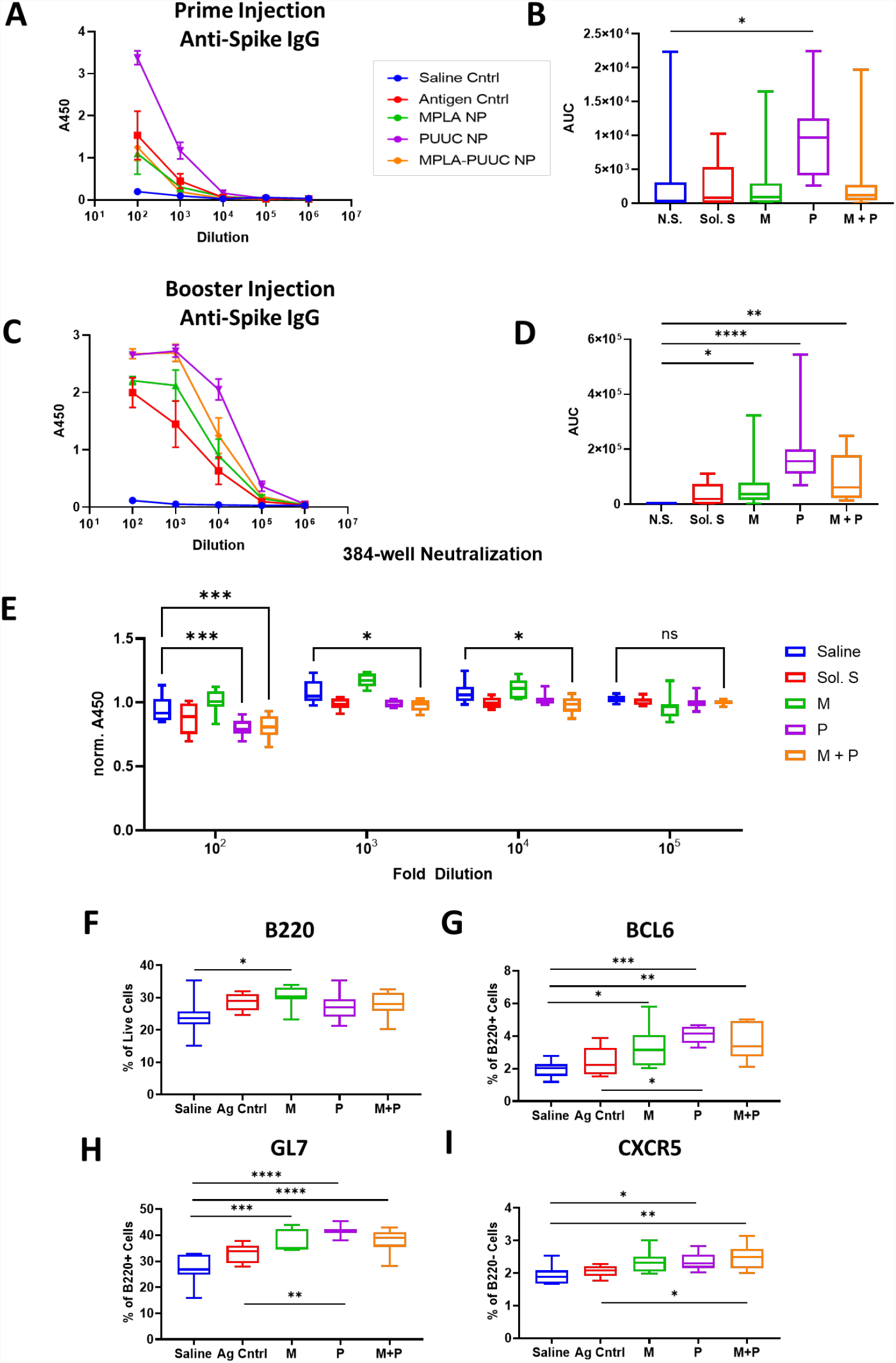
MPLA-PUUC NPs do not increase humoral responses to intramuscular spike protein subunit vaccine compared to PUUC NPs. Female BALB/c mice were immunized by intramuscular injection in the bilateral tibialis anterior muscles on day 0 with saline and spike protein (80 ng) with or without adjuvant-NPs (4 mg) loaded with MPLA (24 μg), PUUC (20 μg), or MPLA+PUUC (24 μg, 20 μg). Pre-booster sera was collected on day 26. On day 28, mice were boosted with either saline, spike protein (1000 ng), with adjuvant-NPs. Mice were euthanized on day 36, and post-booster sera and popliteal lymph nodes were collected. **A)** Total anti-spike IgG quantified by absorbance at 450 nm during ELISA assay and **B)** AUC calculated for each dilution curve. **C)** Total anti-spike IgG in post-booster sera and **D)** AUC for each dilution curve. **E)** Neutralizing anti-spike antibodies quantified with absorbance at 450 nm using modified ELISA assay with biotinylated ACE-2 protein. Lower absorbance values indicate higher neutralizing antibody levels. Absorbance values normalized to blank wells in each row of 384 well plate to correct for plate-based effects. **F)** B220+ out of lymph node cells, **G)** Bcl6+ out of B220+ cells, **H)** GL7+ out of B220+ cells and **I)** CXCR5+ out of B220-cells from the combined popliteal lymph nodes. **B**,**D)** Normality was assessed with the Kolmogorov-Smirnov test. Statistical significance was determined with the Kruskal-Wallis test and Dunn’s post-hoc test for multiple comparisons. **E-I)** Statistical significance calculated with One-Way ANOVA and Tukey post-hoc test. *p ≤ 0.05, **p ≤ 0.01, ***p ≤ 0.001, ****p ≤ 0.0001 for all graphs.

In an analogous experiment using NP-S as antigen (1000/1000 ng S protein) instead of soluble protein, MPLA+PUUC NPs did not lead to a significant increase in anti-spike total IgG before or after a booster injection. Indeed, PUUC-NPs were associated with a decrease in IgG before a booster and both PUUC-NPs and MPLA+PUUC-NPs were associated with a decrease after a booster (**Figure S5**).

## DISCUSSION

In response to studies revealing PAMP combinations synergistically and complementarily enhance immunity, we investigated the singular and combined effects of nanoparticle-delivered TLR and RIG-I agonists on cellular and humoral immune responses against SARS-CoV-2 spike protein. Our data suggest MPLA plus PUUC has the potential to be an effective adjuvant formulation for SARS-CoV-2 protein subunit vaccines and is supported by evidence that the SARS-CoV-2 virus interacts with both TLR4 and RIG-I during natural infection.^38,39^ In vitro data showed MPLA+PUUC, R848+PUUC, and CpG+PUUC PLGA-PEI NPs stimulate more GM-CSF BMDC proinflammatory cytokine secretion (IL-1β, IL-12p70, IFN-β) than single adjuvant controls, and MPLA NPs and CpG-containing NPs (i.e., CpG, CpG+PUUC) maintain or induce CD8+ T cell proliferation with GM-CSF or FLT3L BMDCs, respectively. GM-CSF BMDCs and FLT3L BMDCs, both widely used for evaluating APC maturation *in vitro*, had contradicting responses to TLR and RIG-I agonists most likely because the cultures are comprised of different APC populations (monocytes, monocyte-derived macrophages, and DCs). Consistent with previous reports, we found that GM-CSF BMDCs were primarily composed of a heterogeneous population of monocytes, monocyte-derived macrophages, monocyte-derived DCs, and neutrophils.^34^ FLT3L BMDCs contained higher percentages of conventional DCs and plasmacytoid DCs.^40^ Interestingly, cells derived from the Ly6C^hi^ monocyte lineage are known to secrete IL-27 *in vivo* in response to subunit vaccination with multiple TLR adjuvants, which may explain higher IL-27 production by GM-CSF BMDCs compared to FLT3L BMDCs.^41^

In GM-CSF BMDC culture, MPLA+PUUC NPs induced significant IL-27 production while single-adjuvant MPLA NPs and PUUC NPs were not as potent. IL-27 is a heterodimer of the EBI3 and p28 subunits,^42^ and TLR4 ligation by MPLA generates EBI3 and some p28.^43^ PUUC signaling through RIG-I induces the production of type I interferons, which strongly promote the transcription of p28 through the binding of IRF-1 to IRSEs.^29,43^ It is plausible that IL-27 synthesis in response to MPLA+PUUC is due to the increase in p28 to PUUC-induced type I interferons combined with the increase in p28 and EBI3 to MPLA signaling. Furthermore, possibly due to IL-27-induced expression of T cell inhibitory receptors on APCs such as PD-L1, LAG-3, CTLA-4, and TIGIT,^44^ MPLA+PUUC NPs and R848+PUUC NPs decreased T cell proliferation with GM-CSF BMDCs compared to controls without adjuvant. Alternatively, MPLA NPs increased CD8+ T cell proliferation when cocultured with both GM-CSF and FLT3L BMDCs. This could be explained by the expression of TLR4-induced costimulatory molecules on APCs in the absence of IL-27-induced T cell inhibitory receptors.^45^

Due to observed increases in proinflammatory cytokine production in vitro, we predicted that MPLA+PUUC NPs, R848+PUUC NPs, and CpG+PUUC NPs would elicit stronger cellular and humoral immune responses *in vivo* to SARS-CoV-2 S protein subunit vaccines compared to single-adjuvanted or non-adjuvanted vaccines. Previous studies have reported delivering antigens and adjuvants in separate PLGA particles can improve or attain the same immune responses compared to co-delivered antigens and adjuvants. Therefore, we opted to deliver antigen and adjuvant separately for these studies.^46,47^ Also, because APCs innately recognize the particulate state of microbes, we tested adjuvants with both NP-conjugated (NP-S) and soluble (S-S) SARS-CoV-2 S protein antigens.^48^ CpG-PUUC and R848-PUUC NP delivered with S-S failed to generate significant increases in percentages of T cells producing IFNγ or TNFα. A previous study showed that intramuscular vaccination with R848-PUUC NPs induced lymphopenia, possibly due to the overproduction of type I IFN.^29^ Studies utilizing CpG 1018 or TLR7/8 agonists in SARS-CoV-2 S protein subunit vaccines combine the TLR agonists with alum, which alone traditionally induces a Th2-type immune response.^49–51^

Intranasal vaccination of mice with MPLA+PUUC NPs plus NP-S protein increased CD44^+^ CD4 T cell populations with IFN-γ and TNF-α responses in lung cells during restimulation with SARS-CoV-2 S protein peptide pools. CD44 is a marker that distinguishes effector and memory T cells from naïve subsets.^52^ Because these CD44^+^ CD4 T cells were enriched for intracellular IFN-γ and TNF-α, our results suggest that these cells were polarized towards a Th1 effector phenotype, which may be essential for controlling SARS-CoV-2 infection in humans.^53–55^ Routhu et al. intranasally administered a SARS-CoV-2 RBD trimer subunit vaccine with alum plus 3M-052 (TLR7/8 agonist) in rhesus macaques and also observed a Th1-biased response, but no detectable antigen-specific CD8+ T cell response.^50^ A preclinical study by Jangra et al. investigating a nanoemulsion +/− RIG-I agonist (IVT DI) adjuvanted SARS-CoV-2 S1 subunit intranasal vaccine also observed increased IFNγ secreted by splenocytes and dLN cells, indicating a Th1 response.^56^ Cell populations bearing tissue-resident memory T cell markers (CD69^+^CD103^+^) also increased. Because these T cells were CD4^−^CD8^−^ we speculate they could be γδ T cells, a subset of T cells enriched in epithelial and mucosal tissues that are activated in an MHC-independent manner^57,58^. Unexpectedly, in post-booster BAL fluid and sera, antigen-specific IgG and IgA were present following intranasal vaccination with MPLA NPs plus S-S, but not with PUUC NPs or MPLA+PUUC NPs. Jangra et al. did observe IgG in the BAL fluid and sera but also administered a 15-μg S1 subunit dose, much higher than our 1-μg S protein dose, which may be necessary to induce humoral responses. The addition of RIG-I agonist to their nanoemulsion adjuvant also did not enhance humoral responses against S1 subunit protein.^56^

Intramuscular vaccination with NP-S or S-S plus adjuvant induced very different antigen-specific immune responses compared to intranasal vaccination. Antigen-specific IgG response in mice intramuscularly immunized with NP-S was minimal; IgG was only detectable at 1:100 dilution post-booster in formulations without adjuvant (Figure S5). Conversely, in intramuscular vaccinations with S-S, PUUC NPs generated significant antigen-specific IgG in sera, pre- and post-booster. Post-booster sera from mice immunized with S-S also contained antibodies neutralizing S protein, meaning ACE-2 binding was reduced; these results are noteworthy because SARS-CoV-2 enters respiratory endothelial cells via endocytosis following ACE-2 binding.^59,60^ We also found that S-S plus PUUC NP was associated with increased GC (GC) BCL6^+^ and GL7^+^ B cells in the dLNs. Because the GC reaction produces high-affinity isotype switched antibodies by both plasma cells and memory B cells,^29,61^ we suspect that the increased GC response associated with PUUC NPs explains the increase in antigen-specific-spike IgG and neutralizing antibodies. This GC reaction may be a response to early innate inflammation associated with PUUC, such as endothelial cell activation which has been reported with RIG-I ligation in viral models,^62–64^ and the APC activation which we have observed in our in vitro studies. Because both IgG1 and IgG2a were increased following intramuscular vaccination with PUUC NPs and S-S, we do not attribute the overall increase in antibodies to either a Th1- or Th2-mediated bias.^65^ This finding contrasts with previous studies that have shown a Th2 bias with RIG-I-based adjuvants following intraperitoneal vaccination with influenza VLPs.^66^

Zhou et al. showed that the inclusion of alum in SARS-CoV S protein subunit vaccines reduced the effective antigen dose tenfold. In mice, the alum-adjuvanted 5-μg S protein subunit vaccine produced twice as many neutralizing antibodies as the nonadjuvanted 50-μg S protein subunit vaccine.^67^ Interestingly in our studies, the 80-ng primer and 1000-ng S protein boost produced similar increases in GC B cell populations compared to the 1000-ng primer and 1000-ng boost, indicating an opportunity for antigen dose sparing with the inclusion of PUUC as an adjuvant. MPLA+PUUC NPs plus S-S reduced antigen-specific IgG following the first injection and performed equally to PUUC NPs following the booster injection. MPLA+PUUC NPs and PUUC NPs increased BCL6^+^ and GL7^+^ B cell populations. MPLA+PUUC NPs did significantly induce more neutralizing antibody activity compared to PUUC NPs or MPLA NPs alone, and increased B220^-^CXCR5^+^ cell populations (Tfh markers).

Compared to trials with soluble antigen, it is interesting that MPLA+PUUC NPs failed to elicit a strong anti-spike IgG response when NP-conjugated spike was used instead. This effect could be explained by a hindered ability of NP-conjugated molecules encounter B cells independently, as particles >200 nm in diameter drain less efficiently to lymphatics.^68^ Additionally, given that particle surface topology alone can trigger innate immune responses, particulate antigen might be sufficiently immunogenic enough that when combined with PUUC or MPLA+PUUC, lymphocytes become anergic, giving a reduced adaptive response.^69^

## CONCLUSION

Our results demonstrate that polymer-NP delivery of the RIG-I agonist PUUC), and the TLR4 agonist MPLA, increases immune responses to SARS-CoV-2 spike protein subunit vaccines compared to non-adjuvanted vaccines. MPLA+PUUC NP also elicited differential cellular and humoral responses against SARS-CoV-2 spike protein depending on the APCs encountered (GM-CSF versus FLT3L-derived), route of administration (intramuscular versus intranasal), and delivery-platform of the S protein (soluble versus NP-conjugated). We show that I.N, administration with PUUC+MPLA NPs induces memory T cell responses in the lung while MPLA NPs induces local IgA and IgG responses in the lung. In contrast, I.M. administration induces robust systemic total and neutralizing antibody responses against the SARS-CoV-2 spike protein, Future investigations should examine whether a combination of I.N. and I.M vaccination can produce balanced systemic and lung-specific protective immunity against SARS-CoV-2 challenge and improve vaccine durability and protection against infection and transmission.

## ACKNOWLEDGMENTS

We wish to acknowledge the core facilities at the Parker H. Petit Institute for Bioengineering and Bioscience at the Georgia Institute of Technology for the use of their shared equipment, services, and expertise. These facilities include: the Biopolymer Characterization Core for the preparation of PLGA particles, the Engineered Biosystems Building Physiological Research Laboratory for animal experiments, and the Cellular Analysis and Cytometry Core for flow cytometry experiments. Research reported in this publication was supported in part by the Pediatrics/Winship Flow Cytometry Core of Winship Cancer Institute of Emory University, Children’s Healthcare of Atlanta and NIH/NCI under award number P30CA138292. The content is solely the responsibility of the authors and does not necessarily represent the official views of the National Institutes of Health. This work was partially funded by NIH/NIAID grant U01-AI124270-02 to KR, funds from the Georgia Tech Foundation to KR, the National Science Foundation Graduate Research Fellowship to AA, the NIH T32 Cellular and Tissue Engineering training fellowship (NIH grant T32-GM0843) to AA and AB, and the Robert A. Milton Chaired Professorship to KR.

## CONTRIBUTIONS

A.A., B.P., P.P., and K.R. conceptualized the idea and methodology. K.R. acquired the funding. A.A. was the project administrator and supervisor for iso-MLR assays and A.A. and M.K. were joint supervisors for all in vivo studies. P.P. and B.P. helped A.A. and M.K. conduct in vivo vaccinations. B.P. supervised adjuvant formulation and NP synthesis. A.A., M.K., A.B., and B.P. curated the data. A.A. and M.K. conducted formal analyses. A.A., M.K., B.P., A.B., P.P., C.V., R.J., J.H., C.S., L.K., M.A.O., and D.F. harvested and processed blood and tissue samples from in vivo studies for further investigation. A.A., M.K. and B.P. wrote the manuscript. K.R. revised the manuscript. All authors reviewed the final manuscript and submitted comments.

