## Supplemental Data for "Nanoparticle-delivered TLR4 and RIG-I agonists enhance immune response to SARS-CoV-2 subunit vaccine"

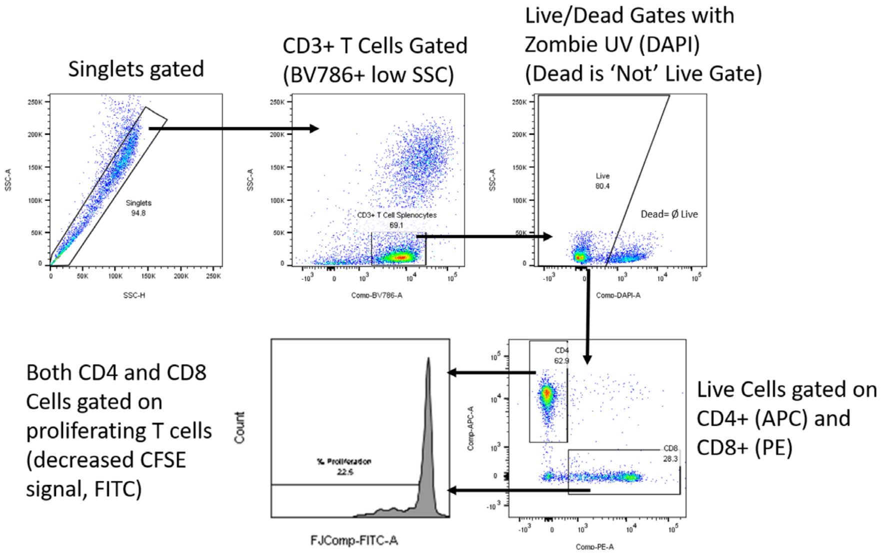


**Supplementary Figure S1. Flow gating scheme for measuring T cell proliferation in iso-MLR assay.**

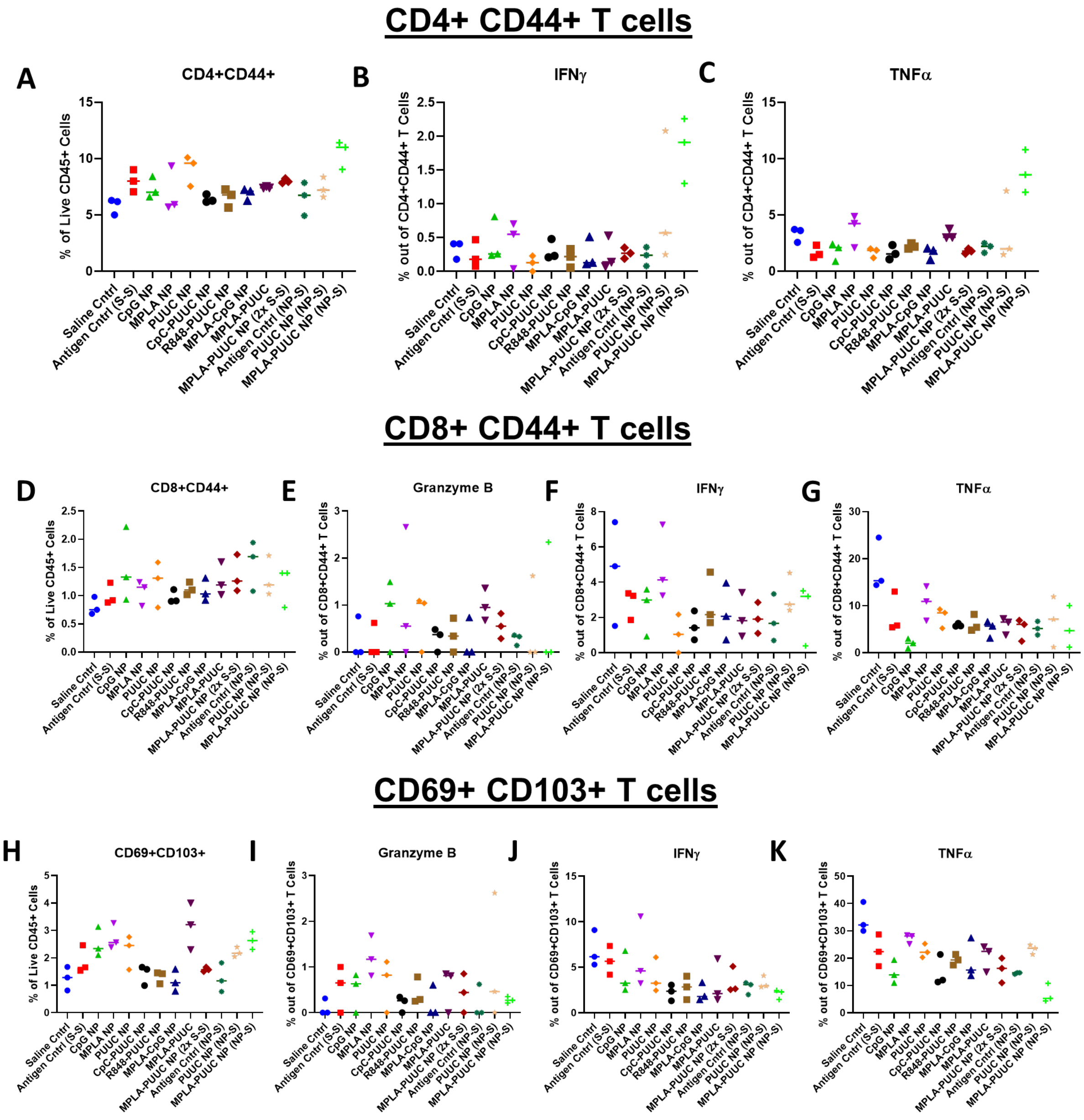


**Supplementary Figure S2. CD4+ CD44+ T cell responses in the lung following intranasal vaccination with combination adjuvants and either soluble or particle-conjugated spike protein.** On days 0 (prime) and 28 (boost), female BALB/c mice were immunized by dropwise addition of saline and unformulated spike protein or spike protein-conjugated NPs (1 µg S protein dose) with adjuvant-NPs (4 mg) (see Materials and Methods for doses). Mice were euthanized and lungs collected on Day 35 (one-week post-boost). Lung cells were restimulated with spike peptide for 6 h. Percentages of cells expressing **A)** CD4+CD44+ out of CD45+ cells, **B-C)** IFNγ+ or TNFα+ out of CD45+CD4+CD44+ cells, **D)** CD8+CD44+ out of CD45+ cells, **E-G)** Granzyme B+, IFNγ+, or TNFα+ out of CD45+CD8+CD44+ cells **H)** CD69+CD103+ (tissue resident memory T cells) out of CD45+ cells, **I-K)** Granzyme B+, IFNγ+, or TNFα+ out of CD45+CD69+CD103+ cells.

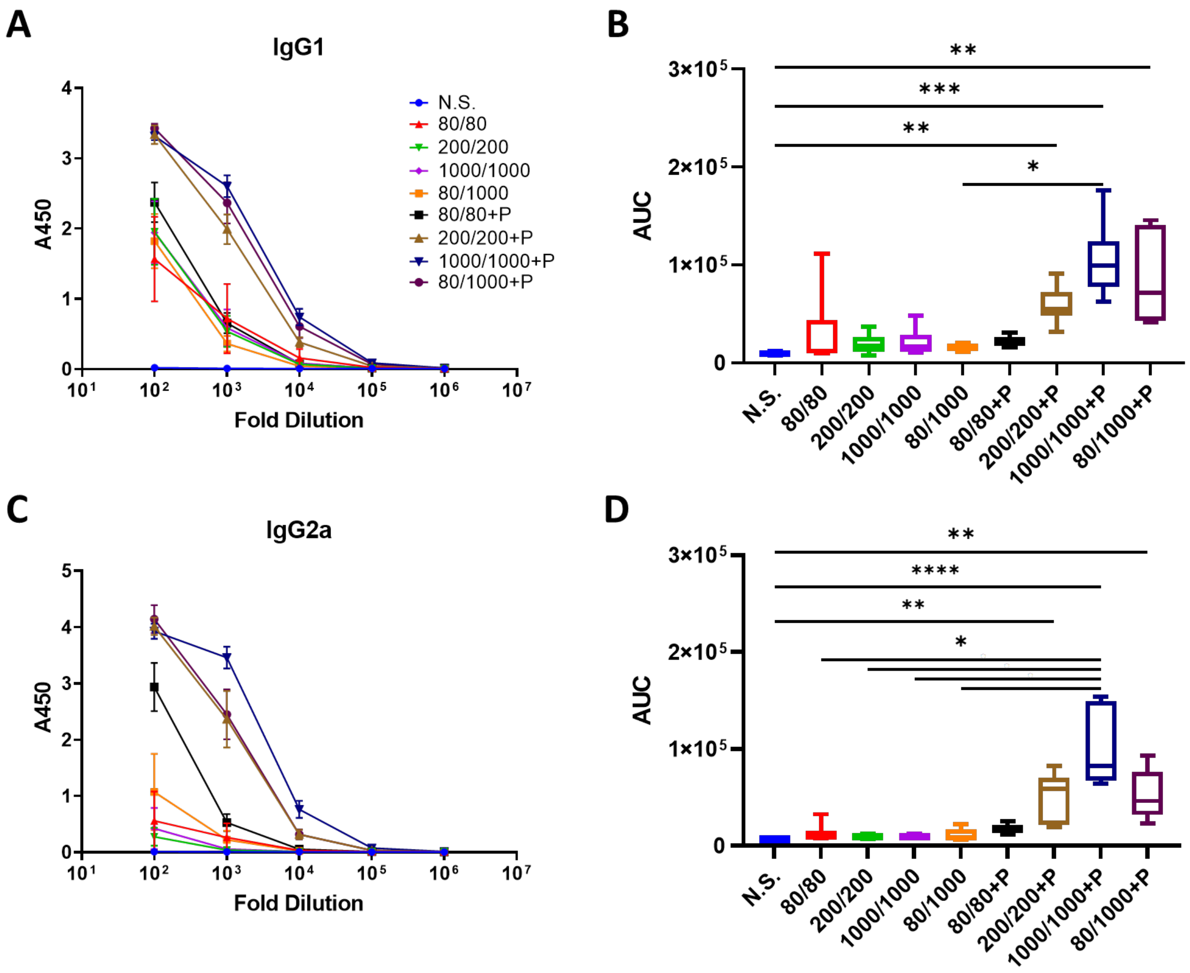


**Supplementary Figure S3: Vaccination with SARS-CoV-2 Spike and PUUC NP adjuvant elicit anti-Spike IgG1 and IgG2a.** Female BALB/c mice were immunized by intramuscular injection into both tibialis anterior muscles at day 0 (prime) with saline or soluble spike protein at 80 ng, 200 ng, 1000 ng with or without adjuvant-NPs (4 mg) loaded with PUUC (20 ng). Peripheral blood was drawn on day 26. On day 28, mice were injected with the same, except that half of the 80 ng prime cohort were given a 1000 ng booster. Mice were euthanized on day 36. **A)** Sera were serially diluted and evaluated for anti-spike IgG1 by ELISA. **B)** Area under the curve (AUC) for each dilution curve was calculated for blood from each mouse. **C)** Anti-spike IgG2a was likewise measured by ELISA and **D)** AUC was calculated and compared for each experimental group. Statistical significance was determined with the Kruskal-Wallis test and Dunn's post-hoc test for multiple comparisons. $^{*}p \leq 0.05$, $^{**}p \leq 0.01$, $^{***}p \leq 0.001$, $^{****}p \leq 0.0001$ for all graphs.

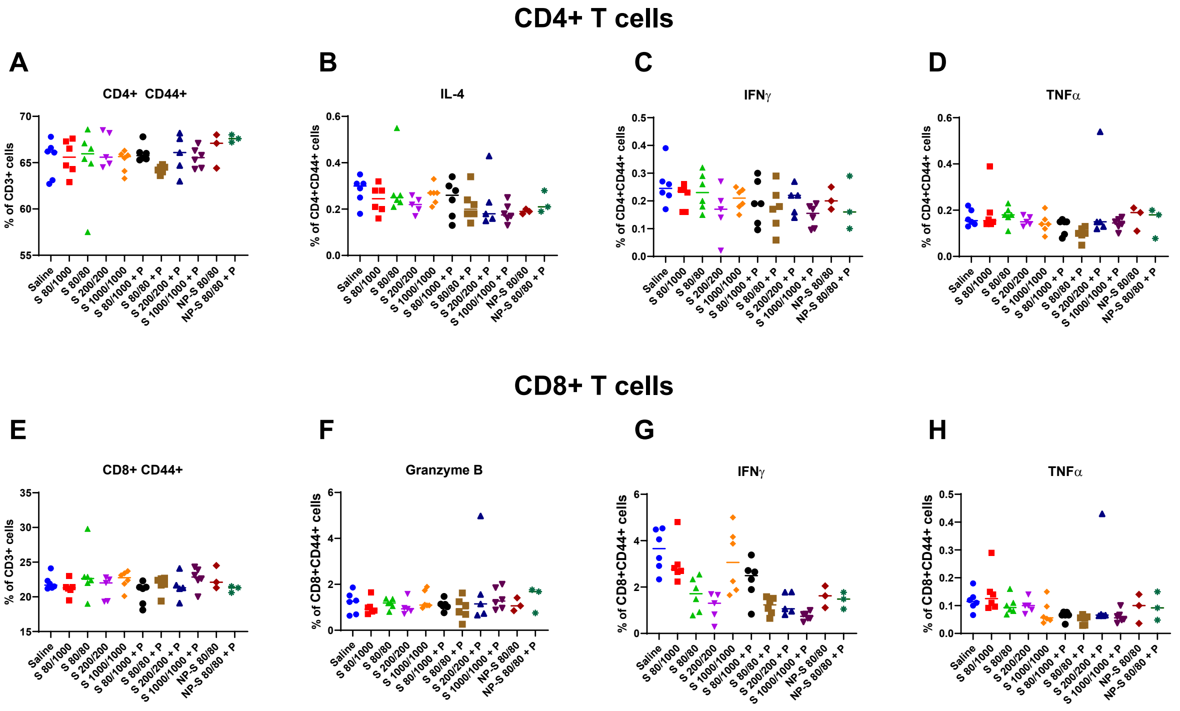


**Supplementary Figure S4. Spleen CD4+ T cell response to restimulation with SARS-CoV-2 spike protein peptides after intramuscular vaccination with PUUC NP adjuvants.** Female BALB/c mice were immunized by intramuscular injection into both tibialis anterior muscles at day 0 (prime) with saline or soluble spike protein at 80 ng, 200 ng, 1000 ng with or without adjuvant-NPs (4 mg) loaded with PUUC (20 µg). On day 28, mice were injected with the same, except that half of the 80 ng prime cohort were given a 1000 ng booster. Mice were euthanized on day 36, and spleens were harvested and plated at 10 million cells/mL (2 million cells/well). Splenocytes were restimulated with SARS-CoV-2 spike peptide pool for 6 h then analyzed by flow cytometry. **A)** Percentage of CD4+CD44+ cells out of CD3+ cells. **B-D)** Percentages of CD4+ CD44+ T cells expressing IL-4, IFN-g and TNF-a. **E)** Percentage of CD8+CD44+ cells out of CD3+ cells **F-H)** Percentages of CD8+CD44+ T cells expressing Granzyme B, IFNγ, and TNFα.

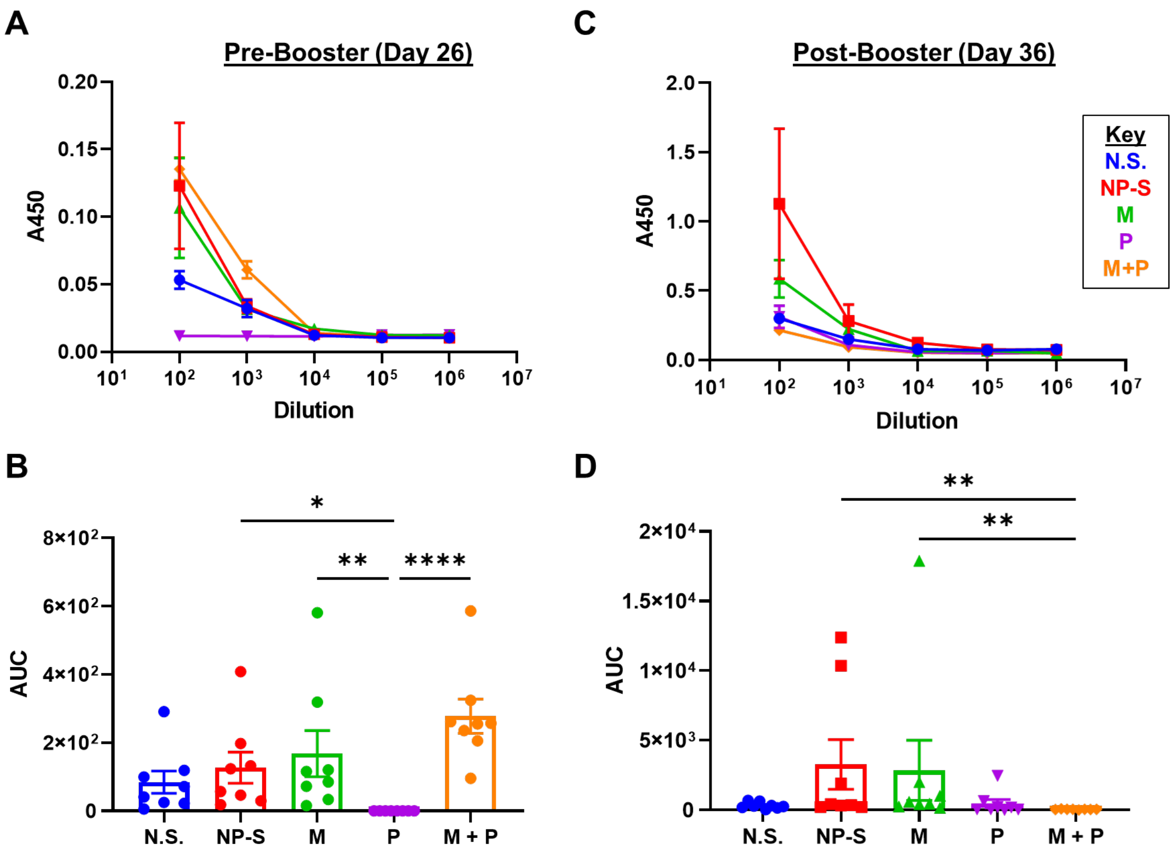


**Supplementary Figure S5. Combination MPLA and PUUC adjuvants fail to precipitate humoral immune responses when delivered with spike protein-conjugated PLGA PEI NPs.** Female BALB/c mice were immunized by intramuscular injection in the bilateral tibialis anterior muscles on day 0 and day 28 with saline and spike-conjugated NPs (NP-S, 1000 ng S protein) with or without adjuvant-NPs (4 mg) loaded with MPLA (24 μg), PUUC (20 ng), or MPLA+PUUC. **A)** Total anti-spike IgG in pre-booster sera quantified using absorbance at 450 nm in ELISA assay and **B)** AUC calculated for each dilution curve. **C)** Total anti-spike IgG in post-booster sera quantified using absorbance at 450 nm in ELISA assay and **D)** AUC calculated for each dilution curve. Absorbances were very low (under 1), a weak signal indicating lack of anti-spike IgG.
